# Three-dimensional tissue stiffness mapping in the mouse embryo supports durotaxis during early limb bud morphogenesis

**DOI:** 10.1101/412072

**Authors:** Min Zhu, Hirotaka Tao, Mohammad Samani, Mengxi Luo, Xian Wang, Sevan Hopyan, Yu Sun

**Author notes:** Authors for correspondence: Sevan Hopyan 686 Bay Street, 16-9713 Toronto, Ontario M5G 0A4 Canada or Yu Sun 5 King’s College Road, MC419 Toronto, Ontario M5S 3G8.

## Abstract

Numerous biophysical hypotheses invoke tissue stiffness as a key parameter for shaping tissue during development and for influencing cell behaviours during disease progression. However, currently available methods are insufficient to test hypotheses that concern the physical properties of bulk tissues. Here we introduce, validate and apply a new 3D magnetic device that generates a uniform magnetic field gradient within a space that is sufficient to accommodate a vertebrate, organ-stage embryo under live conditions. The device allows for rapid, nontoxic measurement of the spatial variation of absolute elastic modulus and viscosity deep within mesenchymal tissues and within epithelia. By applying the device to map the spatiotemporal variation of viscoelastic properties within the early mouse limb bud, we identified an anteriorly biased mesodermal stiffness gradient along which cells move collectively to shape the early bud. Tissue stiffness corresponds to the nascent expression domain of fibronectin that is *Wnt5a*-dependent. The findings challenge the notion that *Wnt5a* regulates cell movements by chemotaxis, and raises the possibility that *Wnt5a* modifies the tissue microenvironment to promote durotaxis *in vivo*. Importantly, the ability to precisely measure tissue stiffness in 3D has the potential to instigate and refine mechanisms of development and disease progression.

## INTRODUCTION

Morphogenesis results from the interrelated effects of tissue scale properties and cellular processes such as neighbour exchange, migration and proliferation. Among epithelia, cellular rearrangements, in part, define tissue scale properties such as anisotropic stress that feedback to orient intercellular rearrangements among invertebrates and vertebrates^1–3^. Far less is understood about mechanisms by which bulk mesenchymal tissues, such as the limb bud, are shaped.

Early limb development represents an accessible model of bulk tissue morphogenesis, and multiple biophysical mechanisms potentially underlie how the limb grows outward from the lateral plate and acquires its particular shape^4^. During early budding of the limb, mesodermal cells move collectively into the limb field from the lateral plate in a *Wnt5a*-dependent manner^5–7^. Based on the movement of mesodermal tissue towards WNT5A-soaked beads that were embedded in the chick embryo, we previously postulated that *Wnt5a* might orient mesodermal cell movements by chemotaxis^5^. However, it remains unclear whether chemotaxis can orient long-range cell movements *in vivo*. *In vitro*, it has long been recognised that cells migrate towards the relatively stiff region of a substrate such as fibronectin, a process termed durotaxis or mechanotaxis^8–10^. Interestingly, increased mesodermal cohesion^11^ coincides with the movement of mesodermal cells into the bud, raising the possibility that tissue properties influence cellular behaviours. However, mechanisms that link these spatial scales have not been sufficiently explored *in vivo*, owing largely to the lack of appropriate tools to map tissue stiffness in three-dimensions within bulk tissues.

Stiffness represents the extent to which an object resists deformation in response to an applied force. Among biological tissues, stiffness varies from a few hundred Pa as in the brain to a few GPa as in cortical bone and is largely dominated by extracellular matrix composition^12^. Several techniques have been employed to measure stiffness in living tissues^13^. Among these, atomic force microscopy (AFM) indentation has been used most extensively to map the surface or epithelial properties of developing plant and animal models^1,14–16^. Although AFM has been applied for deeper mesodermal measurements^14^, it is challenging to decouple surface and deep properties by this method.

Magnetic manipulation devices represent an untethered technique by which magnetic beads or droplets can be displaced or deformed deep inside tissue. Magnetic devices have been employed primarily to measure cellular viscoelastic properties *in vitro*^17^, and in some instances to induce forces upon cells *in vivo*^18–21^. However, few magnetic devices have been employed to determine the spatial distribution of physical properties *in vivo*^22–24^. The application of conventional magnetic devices to quantify tissue mechanical properties *in vivo* is challenging due to low and non-uniform force generation, limited workspace permitted by conventional magnetic devices, and heat generation. Moreover, most available magnetic approaches have been applied in an effectively two dimensional manner, limiting their application to understand the basis of three dimensional cell movements. To circumvent some of these difficulties, a ferrofluid method was developed^22^ which is currently the most suitable technique for measuring mechanical properties deep within tissue. The deformation of injected ferrofluid droplets within the zebrafish tailbud by an array of permanent magnets reflects viscoelastic properties of the tissue. Due to the large size of ferrofluid droplets, usually one droplet is injected per embryo, leading to limited spatial resolution and inability to measure variations and distributions of mechanical properties across tissue.

Additionally, viscoelastic measurements by this approach require 20 minutes due to the slow dynamic response of the droplet, and may reflect the combined influence of material properties and of dissipative processes such as intercellular rearrangements which take place relatively rapidly in zebrafish, *Xenopus* and *Drosophila*.

To quantify tissue stiffness distribution at high spatial resolution, we developed a 3D magnetic device that generates a uniform magnetic field gradient within a workspace that is large enough to accommodate a mouse embryo up to E10.5. The magnetic force generated by the device is sufficient to displace multiple magnetic beads simultaneously to quantify the spatial distribution of stiffness. By applying this device to the mouse embryonic limb bud, we identified a mesodermal stiffness gradient that matches 3D cell movement pattern observed by live light sheet microscopy. The spatial distribution of fibronectin within the early limb field is mediated by *Wnt5a* and mirrors the stiffness gradient we measured. The results support the possibility that durotaxis guides lateral mesodermal cells into the limb.

## RESULTS

### 3D magnetic device for *in vivo* tissue stiffness mapping

Our main design objective was to generate a uniform magnetic field gradient which is crucial for tissue stiffness mapping. In existing magnetic devices, the force exerted on a magnetic bead is large when the bead is close to the magnetic pole tip (which might damage tissue) or is too small to displace the bead when it is far away. In contrast, the magnetic force exerted on a bead within a uniform magnetic field gradient is independent of the distance between the bead and the poles, is identical upon all magnetic beads throughout the field, and does not change as beads move. Other design features of our device included a workspace large enough to accommodate an organ-stage mouse embryo, and the capacity to generate no force perpendicular to the focal plane, thereby ensuring accurate measurement of bead displacements.

The system we created consists of eight magnetic poles with two magnetic yokes and eight coils (**Fig. 1a**). To achieve a uniform magnetic field gradient, the eight poles were arranged into two layers. By vertically aligning the two layers of the poles, a uniform magnetic field gradient can be generated between them^25,26^. The pole-to-pole separation in each layer and the vertical separation of the two layers were carefully chosen to achieve high magnetic field generation and a large workspace to accommodate an entire mouse embryo. The ratio between the vertical separation of the two layers and the pole-to-pole separation was determined by numerical simulation using COMSOL Multiphysics. Among the ratios evaluated (1:0.25, 1:0.5, 1:1, and 1:2), the ratio of 1:0.5 resulted in minimal loss of field strength and field deterioration. The final pole configuration is shown in **Fig. 1b**. The simulated magnetic field of the magnetic device under 2 A driving current is shown in **Supplementary fig. 1a**. The simulation-predicted uniform magnetic field gradient space (**Fig. 1c)** can be approximated as a column with a diameter of 1.2 mm and a height of 1 mm, within which the non-uniformity of the magnetic field gradient is less than 3% (**Supplementary fig. 1b**).

**Figure 1.**
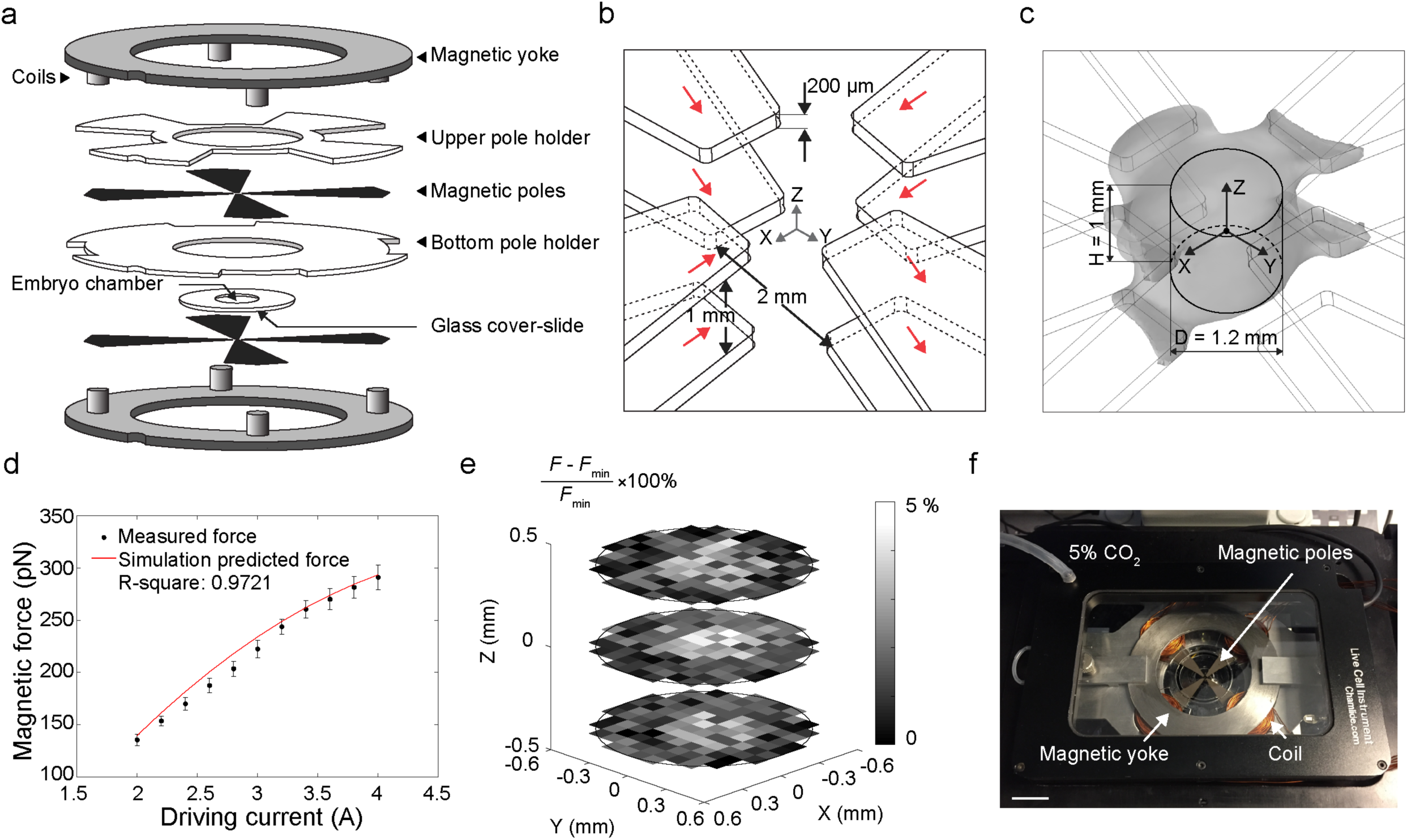
3D magnetic device system. **a**, 3D structure of the magnetic device system. **b**, Configuration of magnetic poles. Red arrow indicates the magnetic flux direction along each pole. **c**, Gray space shows the uniform magnetic field gradient (3% error) predicted by simulation. The uniform magnetic field gradient is approximated as a column (diameter: 1.2 mm, height: 1 mm). **d**, Magnetic force-driving current calibration result. Red line shows simulated force prediction. Error bars represent s.d. **e**, Calibrated force error in three different planes (0.4 mm interval) within the working space. Each grid represents 100 x 100 µm area. **f**, Experimental set-up of magnetic device system on a spinning disc confocal stage with live imaging chamber (scale bar: 2 cm).

We conducted experimental calibration by dispersing 2.8 μm magnetic beads in silicone oils with known viscosities and actuated the magnetic beads to move. The magnetic force was calculated by Stoke’s equation *F*_*drag*_ = 3 *лdηυ* where *d* is the diameter of the bead; is the dynamic viscosity of the silicone oil; v is the velocity of the beads. To ensure the magnetic force exerted on the bead was identical throughout the workspace, each magnetic bead was saturated, according to F= *MV*_*bead*_∇*B*, where *M* is magnetisation and is independent of magnetic field s when saturated; *V*_*bead*_ is the volume of the bead; and vs is the magnetic field gradient. For the magnetic beads selected for use in this work (Dynabead M-280, Invitrogen), the magnetic field needed to saturate the beads must be larger than 0.1 Tesla, which was generated by applying a driving current of at least 2 A as guided by magnetic field simulations (**Supplementary fig. 1a**). The experimentally measured forces agreed well with the simulation-predicted values (**Fig. 1d**) and the deviation of experimentally calibrated forces was less than 5% throughout the workspace, further confirming the uniformity of magnetic field gradient (**Fig. 1e**). We also measured heat generation of our device using a thermocouple probe. Under 10 s continuous actuation with 2 A and 4 A driving currents, the measured temperature change in the workspace centre was less than 0.6 and 1.3 degrees C, respectively (**Supplementary fig. 1c**). The experimentally measured thermal dose under 4 A actuation was 0.01 cumulative equivalent minutes at 43 °C, far below the threshold value of 1 min. for potential thermal damage to tissue^27^. The device was integrated with an incubation chamber and mounted on a spinning disk confocal microscope (**Fig. 1f**).

### Mapping limb bud stiffness

For visualisation under confocal microscopy, the magnetic beads were fluorescently labelled via the streptavidin-biotin reaction. Cell membrane-adhesion molecules were also coupled onto the magnetic beads to ensure that the beads deform cell membranes instead of moving freely within tissue (**Fig. 2a**). Biotinylated fibronectin and biotinylated poly-l-lysine equally prevented unwanted movement of beads upon membranes, and poly-l-lysine was chosen for experiments. The functionalised magnetic beads were microinjected into the limb bud region of mouse embryos. The embryos were labelled with a transgenic membrane marker *mTmG* (membrane-localised tdTomato, membrane EGFP) which was activated to label all cell membranes in the early embryo green using pCX-NLS:Cre^28^. Beads in a volume of 0.5 nL DMEM were deposited at different depths within the 20 somite stage (som., ∼E9.25) mouse limb bud using step-wise microinjection during withdrawal of a glass needle from mesoderm (**Supplementary fig. 2a**). Microinjection of multiple beads did not result in detectable formation of a liquid bubble or tissue separation at confocal resolution, nor in apoptosis following actuation as assessed by immunostaining against caspase 3, an early marker of that process^29^ (**Supplementary fig. 2b**). The majority of the deposited beads (∼93%) successfully adhered to cell membranes, and the distance between two individual beads was consistently over two cell lengths (**Fig. 2b**). Two percent of the deposited beads were intracellular, and 5% were located on a single cell membrane introducing the possibility of inter-bead interaction during actuation (**Supplementary fig. 2c-e**). The latter two situations were excluded from stiffness analysis.

**Figure 2.**
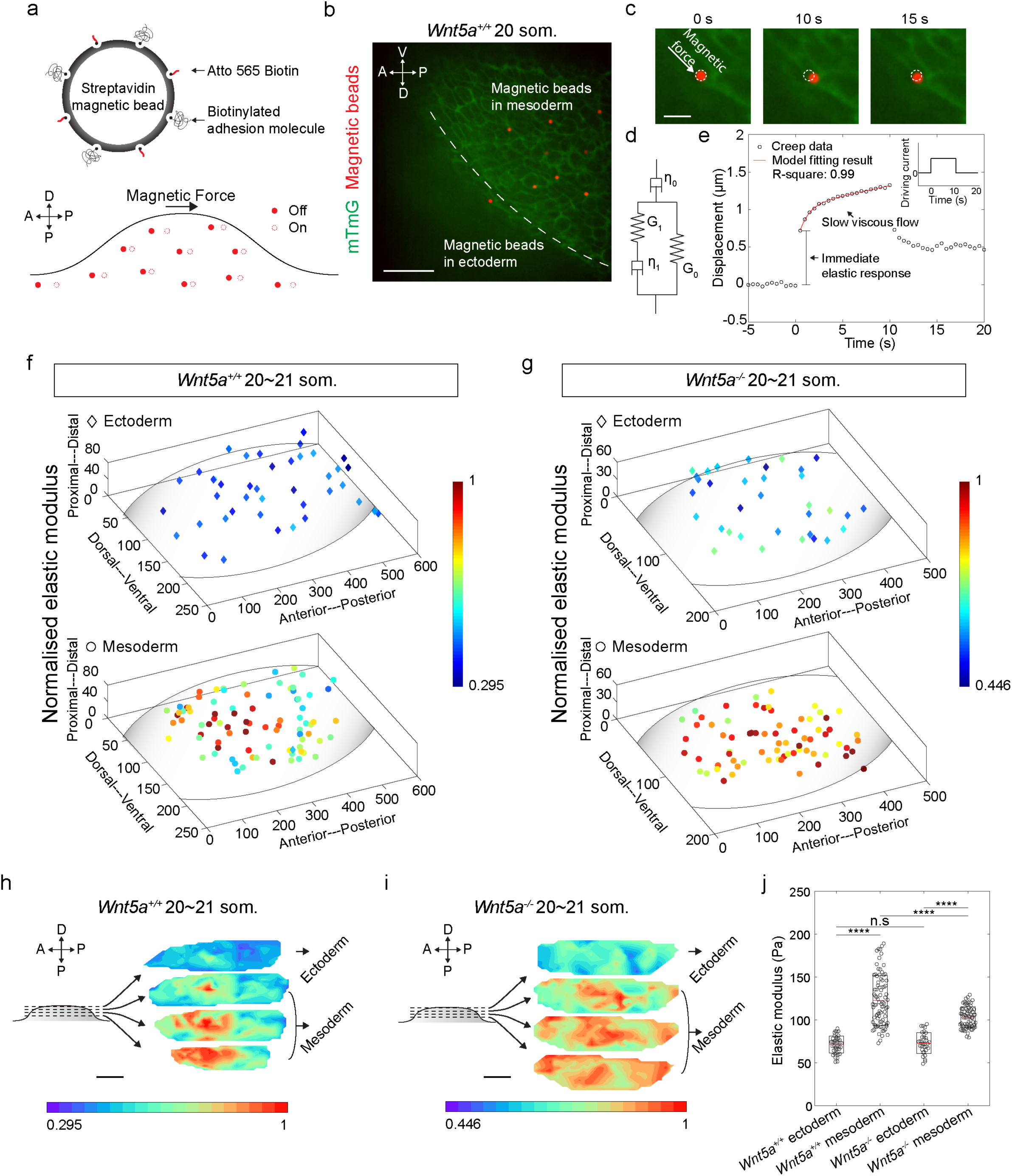
3D tissue stiffness mapping using magnetic device. **a**, Schematic describing magnetic bead functionalisation and actuation during tissue stiffness mapping. **b**, A representative image showing multiple magnetic beads within tissue following step-wise injection. **c**, Higher magnification views of bead displacement during one actuation cycle. **d**, Sketch of standard linear solid model with a serial dashpot diagram characterised by viscous (η_0_ and η _1_) and elastic components (*G*_0_ and *G* _)_. **e**, Representative bead displacement of one actuation cycle characterised by an immediate elastic response followed by a slow viscous flow. Red line shows the model fitting result. Insert shows the driving current during actuation. **f, g**, Normalised elastic modulus (stiffness) map of 20∼21 somite stage WT (*n*=5 embryos) and *Wnt5a*^-/-^ (*n*=4 embryos) limb buds. **h.i**, 3D rendering of the stiffness maps in f and g. **j**, Absolute elastic modulus values of data shown in f and g (two-tailed *t*-test, \*\*\*\**P*<0.0001). Scale bars represent 50 µm (b),10 µm (c) and 100 µm (h, i).

We examined the stiffness distribution within the 20 som. mouse limb bud by actuating the 2.8 μm magnetic beads inside the tissue for 10 s (**Fig. 2c**). Based on the reported speed of mouse limb bud cell migration^5^, cell migration within 10 s should be less than 40 nm, and cell rearrangements take place on substantially longer time scales^1^. Therefore, the effect of morphogenetic movements can be neglected from stiffness measurement. The beads were controlled to deform cell membranes by up to ∼1.5 µm, and bead displacement was continuously measured in the direction of the applied force (**Fig. 2c, e**). Upon actuation, the tissue exhibited an initial elastic response followed by a slow creep response. After the force was removed, the bead retracted partially, not fully, because of the viscoelastic nature of the tissue. We used a standard viscoelastic model (**Fig. 2d, Supplementary fig. 2f** and Methods) to fit the bead displacements (R-square: 0.99). Elastic and viscous components were decoupled using this model. To exclude the stiffness difference between embryos, we normalised the stiffness values by the maximum value for each embryo.

Stiffness mapping revealed that mesoderm of the 20 som. wildtype (WT) limb bud exhibits an anteriorly and proximally biased region of high stiffness that diminishes away from that location. In comparison, the spatial distribution of mesodermal stiffness of 20 som. *Wnt5a*^*-/-*^ mutant embryos was relatively uniform. No stiffness gradient was observed in the ectoderm of either WT or *Wnt5a* mutant embryos (**Fig. 2f-i, Supplementary movie 1, 2**). WT mesoderm exhibited a broader range of values and a higher mean elastic modulus compared to those of the *Wnt5a* mutant (**Fig. 2j**, note: the colour legends in Fig. 2f-i are normalised within each genotype, and do not reflect absolute differences between WT and mutant tissues). The spatial distribution of viscosity closely matched that of stiffness (**Supplementary fig. g-i)**. To validate the ectodermal data measured by the magnetic device, we performed atomic force microscopy (AFM) indentation (**Supplementary fig. 2j**) on the surface of WT embryos. Limb buds were indented with a spherical AFM tip (**Supplementary fig. 2k**) to a shallow depth (<1 µm, **Supplementary fig. 2l**) to mitigate the effect from the underlying mesodermal layer in accordance with the empirical substrate effect rule (indentation depth should be less than 10% sample thickness)^30^. The range of elastic modulus values we measured by AFM agreed closely with those measured using our magnetic device (**Supplementary fig. 2m**). This agreement supports the validity of the magnetic data and that tissue scale properties are properly reflected by the new method.

### Mesodermal cell movement pattern matches stiffness distribution, but does not orient towards *Wnt5a* domain

We previously showed that mesodermal cells enter the early limb field in an anteriorly biased fashion based on live, confocal imaging^5^, suggesting that cell movements may correspond to the spatial distribution of mesodermal stiffness we measured here. To examine this possibility further, we performed live light sheet microscopy of intact *CAG::H2B-GFP* reporter embryos to follow the displacement of cell nuclei in 3D over time. Fluorescent beads were embedded together with the embryo into an agarose gel column that was suspended in media to compensate for positional drift over the course of 2.5 h imaging sessions (**Supplementary movie 3**). Cells were tracked using an autoregressive motion algorithm in Imaris, and drift compensation was performed in Matlab based upon the positions of fluorescent beads embedded alongside the embryo in agarose.

In the WT embryo, cells from lateral plate mesoderm moved into the early limb bud, and ectodermal cells moved toward the midline from dorsal and ventral sides, as expected based on previous confocal time-lapse imaging^1,5^. The 3D nature of the current cell tracking method revealed a previously unrecognised vector of mesodermal and ectodermal cell movement towards an anterior and proximal region of the limb field (**Fig. 3a, Supplementary movie 3, 4**). In the *Wnt5a* mutant, mesodermal and ectodermal cells lacked this convergent pattern of cell movement, and were displaced in a relatively uniform, expansile pattern toward both the anterior and posterior poles of the limb field (**Fig. 3b, c, Supplementary movie 5, 6**). This pattern combined with lower cell migration speeds (**Fig. 3c**) suggests that mutant cell movements may reflect tissue growth rather than collective migration. Together, these findings show that coordinated cell movements correspond to the presence of a gradient in tissue stiffness.

**Figure 3.**
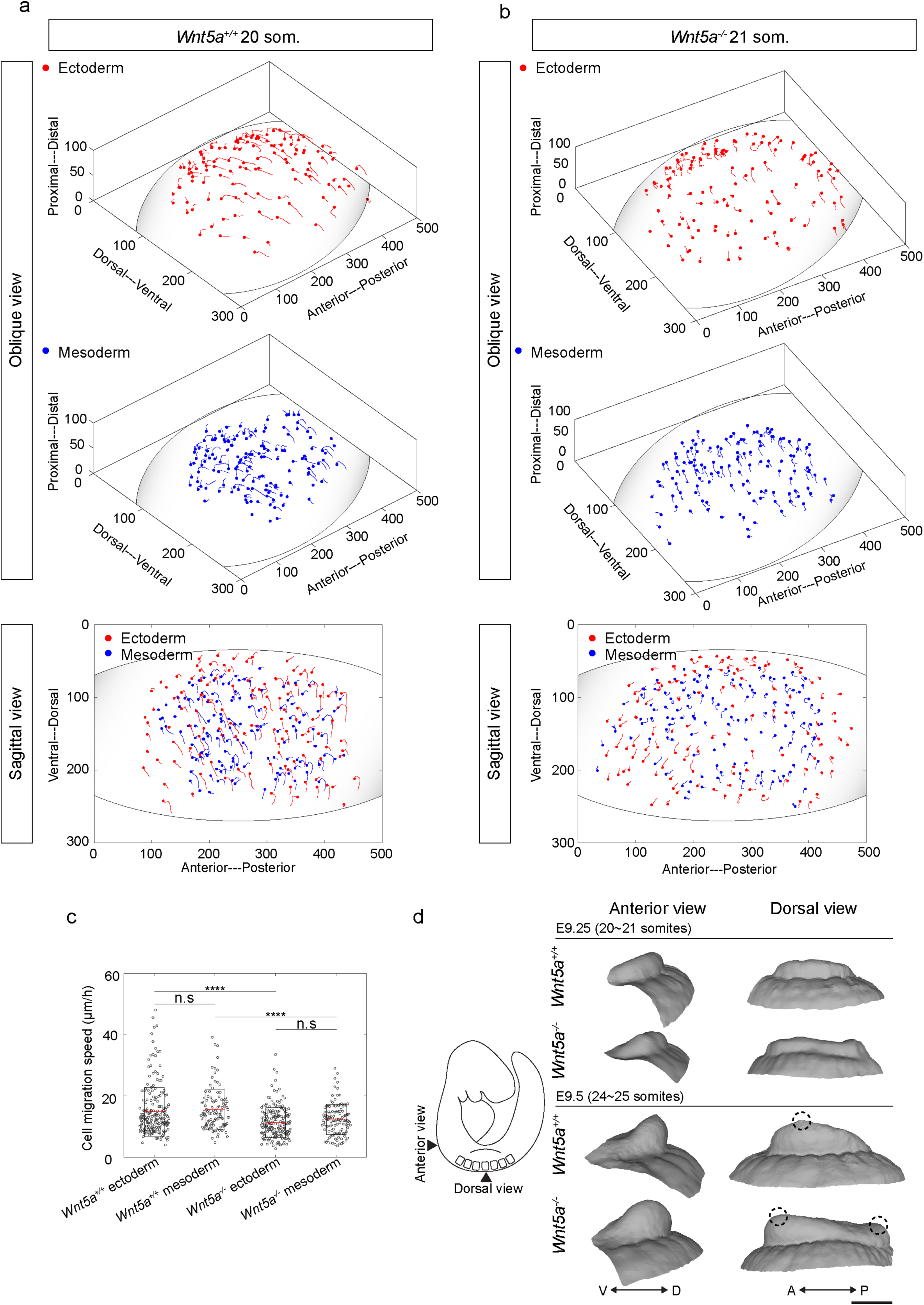
Collective cell migration contributes to early limb bud shape change but is not oriented towards the *Wnt5a* expression domain. **a, b**, 3D cell movement trajectories within 20-21 somite stage WT and *Wnt5a*^-/-^ limb buds tracked by light sheet live imaging (unit: µm). Each dot denotes the last time point of tracking. Grey circle in *Wnt5a*^*+/+*^ mesoderm indicates anteriorly biased region toward which mesodermal cells move. **c**, Cell migration speeds within 20-21 somite WT and *Wnt5a*^-/-^ limb buds. (two-tailed *t*-test, \*\*\*\**P*<0.0001, *n*=3 embryos for each condition) **d**, Limb bud shape change from 20 to 25 somite stage reconstructed from optical projection tomography. Dashed circles indicate the location of distally based peaks of the limb buds. Scale bar represents 200 µm.

The correlation above could be explained by the movement of cells toward a relatively stiff region or to the biased accumulation of cells in a focal region. Our live, membrane-labelled images of the limb field in this study (**Fig. 2b, c, Supplementary fig. 2d**) and in previous studies^1,31^ show that cells are confluent at the resolution afforded by light microscopy. Therefore, packing density can only be changed if cell volume is reduced. Within the relatively stiff, anterior region of the 20 somite stage limb bud, cell density was 170.7 cells/100 µm^3^, and in the posterior region it was 173.3 cells/100 µm^3^, indicating that cell packing does not account for increased anterior stiffness.

To investigate whether tissue is shaped by the morphogenetic cell movements that we observed, we performed optical projection tomography. The WT limb bud acquired an anteriorly biased prominence between 20/21 and 24/25 somite stages (∼E9.25 to E9.5). In contrast, the *Wnt5a* mutant limb bud developed a shallow saddle shape with anterior and posterior prominences (**Fig. 3d**). Our previous work suggested that a saddle-shaped early limb bud is partly attributable to a lack of dorsal and ventral ectodermal convergence^1^, which helps to explain why the early *Wnt5a* mutant bud is not simply bulbous. Of note, immunostaining against phospho-histone H3 (pHH3) and LysoTracker staining suggested that these limb bud shape changes were independent of cell proliferation and apoptosis (**Supplementary fig. 3a-g**). Therefore, in both the WT and mutant cases, tissue shape corresponded to the observed pattern of cell movements.

### Fibronectin distribution in limb bud matches stiffness gradient

*Wnt5a* has been implicated as a directional cue and chemoattractant in the limb bud and other contexts^5,32,33^. Unexpectedly, the 3D vectors of WT cell movements that we observed were not oriented toward the distally biased domain of *Wnt5a* expression^34,35^ (illustrated in **Supplementary fig. 3h**) but rather along a stiffness gradient, raising the possibility that durotaxis underlies collective cell movements in the early limb bud. Since the stiffness gradient is dependent upon *Wnt5a*, downstream factors may play a role in establishing that gradient. Recent *in vitro* work demonstrated that durotaxis depends upon the composition of extracellular matrix (ECM) proteins: cells migrate toward fibronectin-coated, but not laminin-coated, substrate^10^. We examined the spatial distributions of fibronectin and laminin, of cytoskeletal proteins F-actin and vimentin, and of cell-cell adhesion and junctional proteins β-catenin and N-cadherin. Among these candidates, only the expression domain of fibronectin was spatially biased and dependent upon *Wnt5a* (**Fig. 4a-c, Supplementary fig. 4a-m**). Immunostain intensity of fibronectin was strongest in the same anteroproximal region that we identified as the stiffest in the 20/21 som. WT limb bud. Away from that region, fibronectin intensity was attenuated and very weak in the ectoderm and lateral plate. In the absence of *Wnt5a* mesodermal fibronectin was lacking (**Fig. 4a-c**), consistent with findings in the lung where fibronectin is a major target of *Wnt5a* signalling^36^. These results suggest fibronectin establishes a stiffness gradient in a *Wnt5a*-dependent fashion (**Fig. 4d**). The spatial discrepancy in the fibronectin domain and the progressively distal *Wnt5a* domain may be attributable to the time it takes for fibronectin to be synthesised and secreted **(Fig. 4e)**, and anterior bias of the initial fibronectin domain may be due to anterior lateral mesodermal bias of *Wnt5a* expression^34,35^.

**Figure 4.**
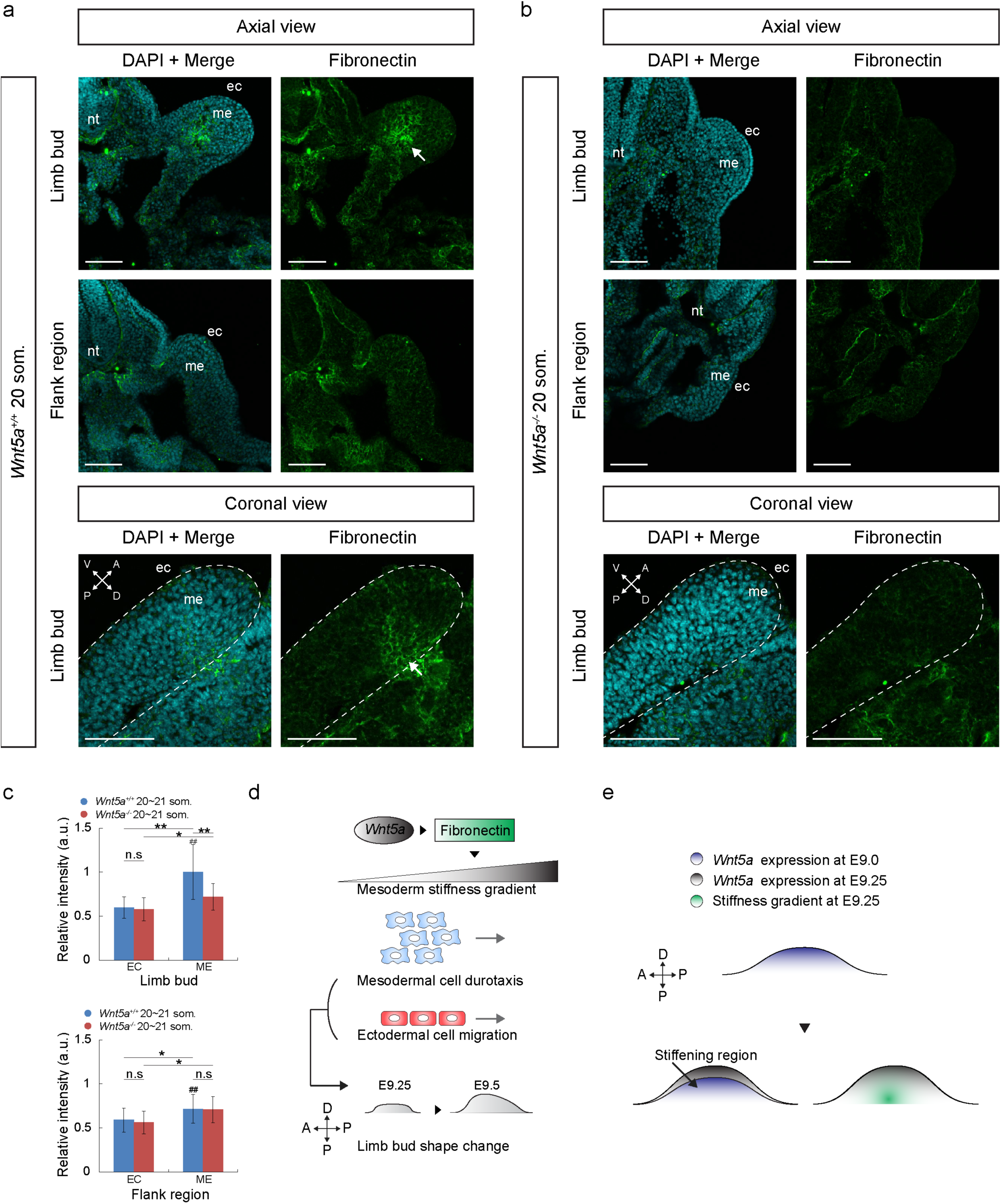
Fibronectin expression domain spatially mirrors the stiffness gradient within the limb bud. **a, b**, Transverse and coronal sections of 20-21 somite WT (a) and *Wnt5a*^-/-^ (b) embryos at forelimb and flank regions. Sections were stained with DAPI (cyan) and anti-fibronectin antibody (green). Arrows indicate the anteriorly biased region of early fibronectin expression. **c**, Relative fibronectin fluorescence intensity of 20-21 somite WT (*n*= 3) and *Wnt5a*^-/-^ (*n*= 3) embryos. **d**, Schematic model representing *Wnt5a* upregulation of fibronectin to generate a stiffness gradient which guides mesodermal cell movements and drives limb bud shape change. Ectodermal cells move in a comparable pattern and may contribute to tissue shape **e**, Graphical description of the potential mechanism by which the stiffness gradient is localised. Error bars indicate s.e.m. Scale bars represent 100 µm in all cases. nt, neural tube; me, mesoderm; ec, ectoderm.

### Stiffness gradient is formed between E9.0 and E9.25

To further examine the putative relationship between fibronectin expression and establishment of a stiffness gradient, we conducted stiffness mapping of 17/18 som. (∼E9.0) embryos when the limb bud first emerges from the lateral plate. At that stage, no stiffness gradient was observed in either WT or *Wnt5a* mutant limb fields (**Supplementary fig. 5a-d, Supplementary movie 7, 8**), and average mesodermal stiffness was lower compared to that of 20/21 som. embryos, particularly in the WT background. Average ectodermal stiffness was not significantly different in either genetic background (**Supplementary fig. 5e**). Corresponding with mesodermal stiffness measurements, fibronectin intensity was spatially uniform and significantly lower than in the 20/21 som. WT embryo (**Supplementary fig. 5f- g**), and collective cell migration was largely absent (**Supplementary fig. 5h-j, Supplementary movie 9- 12**). These findings suggest that tissue stiffness increases following epithelial to mesenchymal transition (EMT) of coelemic epithelial cells that precipitates limb bud initiation^6^. Overall, comparison of data between 17/18 and 20/21 somite stages confirms strong correlation between fibronectin expression, tissue stiffness, collective cell movements and tissue shape.

## DISCUSSION

The resurgence of physical approches to morphogenesis^37^ has been fueled by progressively sophisticated empirical evaluation of mechanical hypotheses. Numerous contemporary proposals in the field invoke feedback between active cell behaviours and tissue scale properties that underlie tissue shape and organisation^2–4,14,22,38,39^. Most of these proposals concern 2D flat or curved tissues for which currently available tools such as microscopic evaluation of cell shape and cytoskeletal organisation, micropipette aspiration, AFM, laser ablation, and ferromagnetic oil droplets have been shown to be sufficient to generate relative or absolute measurements of stiffness, viscosity, cortical tension or stress. However, these tools are not sufficient to assess absolute material properties and their spatial variations within bulk 3D tissues. Emerging mechanisms of morphogenetic phenomena, such as the concept that durotaxis underlies EMT of neural crest cells^14^ and that a viscoelastic gradient contributes to tail bud elongation in zebrafish^22^, have been evaluated using arguably suboptimal methods for measuring bulk tissue properties. These include deep indentation by AFM^14^, which requires potentially imprecise mathematical decoupling of surface and deep layer properties, and ferrofluid approaches that require prolonged periods to assess cell and tissue deformation^22^.

Our magnetic device fills an important void for measurement by generating a uniform magnetic field gradient within a volume sufficient to accommodate an embryo. It is capable of rapidly measuring material properties that are not confounded by morphogenetic cell behaviours such as cell shape changes and rearrangements. Devices similar to the one we introduce may have increasing utility as studies of bulk tissues become more common.

Our application of the magnetic device generated biophysical correlations that can be explained in one of two ways: either morphogenetic cell movements are guided by durotaxis, or cell movements themselves stiffen tissue. Since cell density is not different in the stiffer region, the latter scenario is unlikely. To distinguish formally between these possibilities, spatially precise manipulation of tissue stiffness, ECM proteins, and/or of other potential regulators of ECM will be valuable, and analysis of either potential mechanism in a number of contexts will increase substantially our understanding of how various tissues are shaped during development. The key to opening this line of inquiry has been the newfound capacity to measure previously unattainable parameters with a technological advance.

With regard to the possibility of durotaxis, it is not clear how cells sense a stiffness gradient *in vivo* and potentially titrate active forces to coordinate cell movements. *In vitro*, cells can discern stiffness between 1 Pa/µm^40^ and 400 Pa/µm^41^. The stiffness gradient we measured is ∼0.5 Pa/µm and may reflect finer stiffness sensing ability of cell neighbourhoods *in vivo*. In principle, durotaxis would require the differential binding and unbinding of integrins to specific ECM proteins such as fibronectin. Integrins would engage focal adhesion proteins, such as talin and vinculin, that bind to actomyosin to achieve directional migration towards a stiffer region^42,43^. This process can be mediated by signalling pathways such as non-canonical *Wnt5a* signalling by activating a RhoA-ROCK cascade^44^. Alternatively, cellular forces may also be derived from the differential opening of mechanosensitive ion channels. For example, it has been shown that PIEZO1 can be activated in a stiff environment and inactivated under soft conditions^15^. Experiments to test whether different mechanosensing mechanisms are independent or work cooperatively will be important.

Our technique is readily applicable to other animal models such as zebrafish and *Xenopus laevis*. Beyond developmental biology, the method will also help us to understand the role of stiffness in various diseases processes such as the relationship between tumour stiffness heterogeneity and metastasis, and how increased stiffness is associated with organ (e.g., liver) dysfunction. In addition to the evaluation of tissue scale properties, modification of the methodology we have employed will be applicable to unravelling cellular scale processes such as mechanotransduction. Combining precise physical property measurements with computational simulations of various biological processes will help to generate and test biologically fundamental hypotheses and accelerate discovery.

## METHODS

### Mouse strains

Analysis was performed using the following mouse strains: CAG::H2G-GFP^45^ (Jackson Laboratory: B6.CgTg(HIST1H2BB/EGFP1Pa/J), mTmG^46^ (Jackson Laboratory: Gt(ROSA)26Sortm4(ACTB-tdTomato-EGFP)Luo/J)), pCX-NLS:Cre^28^ (Jackson Laboratory: NMRI.Cg-Tg(CAG-cre)1Nagy/Cnbc), *Wnt5a*^*+/-*35^. All strains were outbred to CD1, and all animal experiments were performed in accordance with protocols approved by the Animal Care Committee of the Hospital for Sick Children Research Institute.

### Optical projection tomography

Mouse embryos were harvested and fixed in 4% paraformaldehyde overnight at 4°C. OPT was performed using a system that was custom-built and is fully described elsewhere^47^. Three-dimensional data sets were reconstructed from auto-fluorescence projection images acquired during a 25 min. scan period at an isotropic voxel size of 4.5 µm. The limb bud structure was segmented from the embryo and rendered in MeshLab.

### Live, time-lapse light sheet microscopy

Three-dimensional time-lapse microscopy was performed on a Zeiss Lightsheet Z.1 microscope. Embryos were suspended in a solution of DMEM without phenol red containing 12.5% filtered rat serum, 1% low-melt agarose (Invitrogen) and 2% fluorescent beads (SigmaAldrich, 1:500, diameter: 500 nm) that were used for drift-compensation, in a glass capillary tube. Once the agarose solidified, the capillary was submerged into an imaging chamber containing DMEM without phenol red, and the agarose plug was partially extruded from the glass capillary tube until the portion containing the embryo was completely outside of the capillary. The temperature of the imaging chamber was maintained at 37° C with 5% CO_2_ Images were acquired using a 20 X/0.7 objective with dual-side illumination, and a *z*-interval of 0.5 µm. Images were acquired for 2-3 hours with 10 min. intervals.

### *In vivo* drift-compensated cell tracking

The light sheet time-lapse image was first rendered in Imaris (Bitplane). The positions of cell nuclei were tracked over time using an autoregressive motion algorithm. Ectodermal and mesodermal cells were separated based on mean thresholding of fluorescence intensity. The tracking data were then imported into R2017b Matlab (MathWorks) for drift compensation using a customised program.

### Magnetic device

The uniform magnetic field gradient was generated by an octuple-pole magnetic device. In order to generate a large magnetic field gradient, the magnetic poles were made of silicon iron (MµShield) which has high permeability at high induction and high saturation. Each pole piece was fabricated from a silicon iron sheet 200 µm thick by electrical discharge machining. The magnetic yoke and the stage were machined by computer numerical control (CNC) from low carbon STEEL 1018 and aluminum, respectively. Each core was wound 100 times with American wire gauge 24 copper wire. Holders for the magnetic poles were cut from an acrylic sheet 1 mm thick by laser machining. Magnetic poles were aligned under the microscope using a calibration slide (MR400, AmScope). The magnetic device was mounted on a Quorum Wave FX-X1 spinning disk confocal system (Quorum Technologies Inc.) that supports a live imaging chamber.

### Magnetic force calibration

To calibrate the magnetic force generated by our magnetic device, magnetic beads (Dynabead M-280, Invitrogen) were dispersed in silicone oil (SigmaAldrich) of known viscosity. In detail, 5 µl of beads and 1 mL of silicone oil were placed in an eppendorf tube. To avoid bead aggregation, ultrasound (Model 60, Fisher Scientific) was used to fully mix the solution. The bead-silicone oil solution was placed in the imaging chamber. After a few minutes when all flows in the solution were settled, the device was activated and the bead movements were recorded (12 fps). Bead velocities were calculated in ImageJ using the Particle Detector & Tracker plug-in. Force exerted on the beads was calculated using the Stokes drag equation F = 3*лdηυ*.

### Magnetic field simulation

Magnetic field simulation of the device was performed in COMSOL Mutiphysics 5.3 (COMSOL Inc.). The HB (magnetisation) curves of silicon iron and LC STEEL 1018 were imported into the software and assigned to pole and yoke, respectively. A sweep function (from 2 A to 4 A with 0.2 A step size) was incorporated in the simulation to derive the magnetic field under different driving currents. Partial differential equations were applied to calculate the spatial derivative of the magnetic field (i.e., magnetic field gradient). Within the workspace, simulation predicted that the non-uniformity of the magnetic field gradient was less than 3%.

### Magnetic bead functionalisation and microinjection

The streptavidin-coated Dynabead M-280 superparamagnetic beads were coupled with Atto565 biotin (SigmaAldrich) and biotinylated adhesion molecules through the streptavidin biotin reaction. We placed 5 μL streptavidin-coated magnetic bead solution into an eppendorf tube. The bead solution was washed three times with phosphate-buffered saline (PBS) to remove preservatives. A permanent magnet was placed under the tube to collect the magnetic beads (i.e., magnetic separation). The beads were then collected by a micropipette and re-suspended in 90 μL of Milli-Q. Atto565 biotin (1 mg) was diluted in 200 μL of ethanol. Five μL of this dilution and 5μL of the biotinylated adhesion molecule solution (1:1,000 in PBS) were mixed with 90 μL of the magnetic bead suspension for 30 min.. Finally, the solution was washed with PBS five times to remove the biotin surplus through magnetic separation, and the supernatant was collected using a micropipette.

For microinjection, 5 μL of functionalised magnetic bead solution was suspended in 100 μL DMEM. The solution was then loaded into a microneedle pulled from a glass capillary tube using a micropipette laser puller. A microinjector (CellTram 4r Oil, Eppendorf) was used to inject magnetic beads into the limb bud at anterior, middle and posterior regions. Multiple depositions were made during one penetration and withdrawal of the needle to distribute the magnetic beads at different depths within the limb bud.

### Stiffness mapping *in vivo*

The embryo with magnetic beads injected was placed into the customised imaging chamber and immobilised by DMEM without phenol red containing 12.5% rat serum and 1% low-melt agarose (Invitrogen). The temperature of the imaging chamber was maintained at 37° C with 5% CO_2_. Prior to stiffness mapping, a Z scan was taken to record the bead locations within the limb bud. Magnetic beads were then actuated by the magnetic device and bead displacements were recorded by spinning disk confocal microscopy at the highest frame rate. Displacement of beads in the direction of the magnetic force was tracked in ImageJ using the Particle Detector & Tracker plug-in. Bead displacements were fit using a viscoelastic model in Matlab to decouple the elastic and viscous components.

Different viscoelastic models have been applied in the literature, for example, Power law^39^, generalised Kelvin^48^ and generalised Maxwell^22^. The relation between creep compliance *J* and time under a step stress varies between these models. Based on our measurements, tissue exhibited a clear initial elastic response upon the application of force. Compared with the Power law and generalised Kelvin models, the generalised Maxwell model was better able to capture the initially elastic response. However, the generalised Maxwell model predicts a plateau at later time points while our data showed continued deformation (**Supplementary fig. 2f**). To fully capture the viscoelastic behaviour we observed, a serial dashpot was added to a standard linear solid model to account for continued deformation. The creep compliance J of the model is

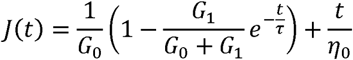

The displacement of the bead is related to the force applied

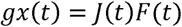

where *g* is a geometric factor. For beads embedded in the material, *g* = 3*лd* with *d* being the diameter of the bead. The shear modulus was converted to elastic modulus *E* by

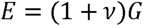

where Poisson’s ratio *ν* = 0.4 (ref.^49^).

The stiffness map was a set of points in 3D, each mapped to their corresponding normalised elastic modulus. Using a standard Delaunay triangulation technique, a set of tetrahedrons was obtained with these points positioned at the vertices. The value of the elastic modulus was approximated inside each tetrahedron by calculating the barycentric coordinates. The result was a continuous field for normalised elastic modulus. Section cuts were made perpendicular to the proximal-distal axis at 80%, 60%, 40% and 20% of the total length as depicted in **Fig. 2h, i** and **Supplementary fig. 5c, d**.

### Atomic force microscopy indentation

Mouse embryos were incubated in 50% rat serum and DMEM in a 35 mm dish in which 2% agarose was poured around the perimeter. The embryo was pinned to the agarose with pulled glass needles placed well away from the limb field. The limb bud was then examined using an AFM (BioScope Catalyst, Bruker) mounted on an inverted microscope (Nikon Eclipse-Ti). AFM indentation tests were performed using a spherical tip, 35 µm in diameter (**Supplementary Fig. 2k**). Spherical tips were made by assembling a borosilicate glass microsphere onto a tipless AFM cantilever using epoxy glue. The cantilever spring constant was calibrated each time before experimentation by measuring the power spectral density of thermal noise fluctuation of the cantilever under no load. To determine the elastic modulus of ectoderm, a trigger force of 250 pN was consistently applied. Because ectodermal thickness was approximately 10 µm, we followed the empirical 10% rule to indent the surface up to 1 µm in depth to avoid the influence from the underlying mesoderm. The Hertz model for the spherical tip was used to calculate ectodermal elastic modulus from the experimental indentation data.

### Immunofluorescence

Dissected mouse embryos were fixed overnight in 4% paraformaldehyde in PBS followed by 3 washes in PBS. Fixed embryos were embedded in 7.5% gelatin / 15% sucrose and sectioned into 10 µm slices using a Leica CM1800 cryostat. Sections were washed 2 x 5 min in Milli-Q and 1 x 5 min in PBS, permeabilised in 0.1% Triton X-100 in PBS for 20 min. and blocked in 5% normal donkey serum (in 0.05% Triton X-100 in PBS) for 1 h. Sections were incubated in primary antibody overnight at 4°C followed by 4 x 10 min. washes in 0.05% Triton X-100 in PBS, then incubated in secondary antibody (1:1000) for 1 h at room temperature. Finally, sections were washed 3 x 5 min in 0.05% Triton X-100 in PBS and 2 x 5 min. in PBS. Images were acquired using a Nikon A1R Si Point Scanning Confocal microscope at 10X, 20X or 40X magnification, and analysis was performed using ImageJ.

### Cell apoptosis detection

Lyso Tracker Red DND-99 (ThemoFisher) was diluted to 2 µM in DMEM containing 50% rat serum. Embryos were placed in the medium and incubated in a roller culture apparatus for 1 h. The temperature was maintained at 37° C with 5% CO_2_. Embryos were washed three times with PBS after staining to remove Lyso Tracker surplus, then fixed overnight in 4% paraformaldehyde in PBS followed by 3 washes in PBS. Images were acquired using a Nikon A1R Si Point Scanning Confocal microscope at 20X magnification, and analysis was performed using ImageJ.

### Antibodies

Laminin (Sigma, 1:100); fibronectin (Abcam,1:100); vimentin (Sigma, 1:500); N-cadherin (BD Bioscience, 1:250); β-catenin (BD Bioscience, 1:1,000); phospho-histone H3 (Cell Signaling, 1:250); caspase 3 (Cell signaling, 1:200); Rhodamine phalloidin (Invitrogen, 1:1,000). All secondary antibodies were purchased from Jackson Immunoresearch and used at 1:1,000 dilutions.

## ACKNOWLEDGEMENTS

We thank Rudolph Winklbauer for critically reviewing the manuscript. This work was funded by the Canadian Institutes of Health Research to SH (MOP 126115) and the Natural Sciences and Engineering Research Council of Canada (NSERC) through a Discovery Grant to YS (RGPIN-2018-06061). YS also acknowledges financial support from the Canada Research Chairs Program.

## SUPPLEMENTARY FIGURE LEGENDS

**Supplementary figure 1. 3D magnetic field within the device. a**, Simulated magnetic field under 2 A actuation plotted in X-Y, X-Z and Y-Z planes as shown in Fig. 2c. The dashed line shows the boundary of the working space. Arrow indicates the minimum magnetic field in the working space. **b**, Simulated magnetic field gradient distribution in three different planes within the working space. **c**, Temperature change in the working space centre under 10 s of 2 A and 4 A actuation. Error bar indicates s.d..

**Supplementary figure 2. 3D tissue stiffness mapping using magnetic device. a**, Step-wise microinjection of magnetic beads during retraction of the glass needle from the limb bud. Asterisks indicate the needle tip position. **b**, Confocal image of injected mouse limb bud visualising DAPI (cyan), magnetic beads (red) and anti-caspase 3 immunostain (green). **c**, Illustration of three distinct bead positions that were observed following microinjection. Case 1: deposited bead successfully adhered to a cell membrane either in between (a) or inside cells (b). Case 2: more than one bead adhered to a single cell membrane. Case 3: bead trapped inside a cell. **d**, Representative images of different bead positions shown in c. **e**, Pie graph shows the percentage of different bead positions. **f**, Viscoelastic models that have been applied previously to quantify tissue properties and their creep compliance-time relations. The fourth example on the right most fully captured the viscoelastic properties that we observed. **g**, **h**, Normalised viscosity map of 20∼21 somite stage WT (*n*=5 embryos) and *Wnt5a*^-/-^ (*n*=4 embryos) limb buds. **i**, Absolute viscosity values of data shown in g and h (two-tailed *t*-test, \*\*\*\**P*<0.0001). **j**, Schematic represents AFM indentation of mouse limb bud. **k**, Scanning electron micrograph of an AFM cantilever with a spherical tip of 35 µm diameter. **l**, A representative force-displacement curve under small indentation (indentation depth <1 µm). **m**, Comparison of ectodermal elastic modulus measured using AFM or our magnetic device. Scale bars represent 100 µm (a, b) and 10 µm (d).

**Supplementary figure 3. Mouse limb bud shape change is independent of cell proliferation and apoptosis. a**, Confocal image of 20 somite WT and *Wnt5a* mutant limb buds visualising DAPI (cyan) and phospho-histone H3 (red). **b**-**e**, Histogram representing percentage of pHH3-positive cells in WT (ectoderm (b); mesoderm (c)) and *Wnt5a* mutant (ectoderm (d); mesoderm (e)) limb bud. **f, g**, Confocal image of 20 somite WT and *Wnt5a*^-/-^ limb buds visualising LysoTracker (red). Scale bars represent 100 µm. **h**, Schematic representation of the *Wnt5a* expression domain.

**Supplementary figure 4. Immunostaining against ECM, cytoskeletal and cell-cell adhesion proteins within the limb field. a**-**d**, Transverse sections of 20 somite WT (a, c) and *Wnt5a*^-/-^ (b, d) forelimb and flank regions. Sections were stained with DAPI (cyan), anti-laminin antibody (green) and anti-vimentin antibody (red). **e**-**m**, Confocal sections of 20 somite WT (e, f, i, j) and *Wnt5a*^-/-^ (g, h, k, m) limb buds visualising DAPI (cyan), F-actin (red), anti-β-catenin antibody (green in e-h) and anti-N-cadherin antibody (green in i-m). *n*=3 embryos for each condition. Scale bars represent 100 µm in all cases. nt, neural tube; me, mesoderm; ec, ectoderm.

**Supplementary figure 5. Stiffness gradient is absent and cell migration is diminished in 17-18 somite stage limb buds. a, b**, Normalised elastic modulus (stiffness) maps of 17-18 somite WT (*n*=3 embryos) and *Wnt5a*^-/-^ (*n*=3 embryos) limb buds. **c, d**, 3D rendering of the stiffness maps of a and b. **e**, Absolute elastic modulus values of data shown in a and b (two-tailed *t*-test, \*\*\*\**P*<0.0001). **f**, Transverse sections of 17 somite WT and *Wnt5a*^-/-^ limb buds. Sections were stained with DAPI (cyan) and anti-fibronectin antibody (green). **g**, Relative fluorescence intensity of fibronectin in 17 somite WT (*n*= 3 embryos) and *Wnt5a*^-/-^ (*n*= 3 embryos) limb fields. **h, i**, 3D cell trajectories within 17 somite WT and *Wnt5a*^-/-^ limb fields tracked from live light sheet time lapse movies (unit: µm). Each dot denotes the last time point of tracking. **j**, Cell migration speed of 17 somite WT (*n*=3 embryos) and *Wnt5a*^-/-^ (*n*=3 embryos) limb buds. Scale bars represent 100 µm. nt, neural tube; me, mesoderm; ec, ectoderm.

## MOVIE LEGENDS

**MOVIE 1** Three dimensional normalised stiffness distribution within 20-21 somite stage WT limb bud. Circles and diamonds represent mesodermal and ectodermal measurements, respectively. Position 0 of the AP, DV and PD axes reflects anterior, dorsal and proximal respectively.

**MOVIE 2** Three dimensional normalised stiffness distribution of 20-21 somite stage *Wnt5a*^-/-^ limb bud. Circles and diamonds represent mesodermal and ectodermal measurements, respectively. Position 0 of the AP, DV and PD axes reflects anterior, dorsal and proximal respectively.

**MOVIE 3** Three dimensional rendering and time lapse movie of a 20 somite *CAG::H2B-GFP* (WT) transgenic embryonic limb bud imaged by light sheet microscopy. Anterior is to the left, dorsal is upwards. The counter in the lower right denotes 12 time steps at 10 min. intervals.

**MOVIE 4** Three dimensional migration trajectories of a subset of cells of a 20 somite *CAG::H2B-GFP* (WT) transgenic embryonic limb bud imaged by light sheet microscopy. Blue and red tracks represent mesodermal and ectodermal cell movements, respectively. Position 0 of the AP, DV and PD axes reflects anterior, dorsal and proximal respectively.

**MOVIE 5** Three dimensional rendering and time lapse movie of a 21 somite *CAG::H2B-GFP;Wnt5a*^-/-^ transgenic embryonic limb bud imaged by light sheet microscopy. Anterior is to the left, dorsal is upwards. The limb bud is in the centre (not the brighter region in the lower left of the frame).

**MOVIE 6** Three dimensional migration trajectories of a subset of cells of a 21 somite *CAG::H2B-GFP;Wnt5a*^-/-^ transgenic embryonic limb bud imaged by light sheet microscopy. Blue and red tracks represent mesodermal and ectodermal cell movements, respectively. Position 0 of the AP, DV and PD axes reflects anterior, dorsal and proximal respectively.

**MOVIE 7** Three dimensional normalised stiffness distribution within 17 somite stage WT limb bud. Circles and diamonds represent mesodermal and ectodermal measurements, respectively. Position 0 of the AP, DV and PD axes reflects anterior, dorsal and proximal respectively.

**MOVIE 8** Three dimensional normalised stiffness distribution of 17 somite stage *Wnt5a*^-/-^ limb bud. Circles and diamonds represent mesodermal and ectodermal measurements, respectively. Position 0 of the AP, DV and PD axes reflects anterior, dorsal and proximal respectively.

**MOVIE 9** Three dimensional rendering and time lapse movie of a 17 somite *CAG::H2B-GFP* (WT) transgenic embryonic limb bud imaged by light sheet microscopy. Anterior is to the left, dorsal is upwards.

**MOVIE 10** Three dimensional migration trajectories of a subset of cells of a 17 somite *CAG::H2B-GFP* (WT) transgenic embryonic limb bud imaged by light sheet microscopy. Blue and red tracks represent mesodermal and ectodermal cell movements, respectively. Position 0 of the AP, DV and PD axes reflects anterior, dorsal and proximal respectively.

**MOVIE 11** Three dimensional rendering and time lapse movie of a 17 somite *CAG::H2B-GFP;Wnt5a*^-/-^ transgenic embryonic limb bud imaged by light sheet microscopy. Anterior is toward the upper left, dorsal is towards the upper right. The limb field is in the central, relatively dark part of the frame.

**MOVIE 12** Three dimensional migration trajectories of a subset of cells of a 17 somite *CAG::H2B-GFP;Wnt5a*^-/-^ transgenic embryonic limb bud imaged by light sheet microscopy. Blue and red tracks represent mesodermal and ectodermal cell movements, respectively. Position 0 of the AP, DV and PD axes reflects anterior, dorsal and proximal respectively.

